# Reinforcement learning enables single-cell foundation models to learn cellular differentiation

**DOI:** 10.64898/2025.12.09.693267

**Authors:** Kai-Li Chang, Hu Chen, Zhandong Liu

## Abstract

While single-cell foundation models excel at static representation learning and single-step perturbation prediction, their capacity to model and control dynamic, sequential cell state transitions remains underexplored. Here, we introduce Differentiation with Reinforcement Learning (DiRL), a framework that transforms foundation models into predictive environments for optimizing multi-step perturbation strategies. By formulating differentiation as a goal-conditioned sequential decision-making problem, DiRL trains agents to navigate the high-dimensional latent landscape of gene expression, learning policies that direct stem cells toward specific terminal fates. Evaluated on iPSC-derived organoid datasets, DiRL outperforms random perturbation baselines with a 69% win rate, with performance scaling systematically as the planning horizon increases. Beyond optimization, DiRL offers interpretable insights into the mechanics of cell fate transitions: the learned policies successfully recover known sequential regulators in Wnt and Hedgehog signaling pathways, while the value function from the critic model recapitulates biological pseudotime orderings comparable to established trajectory inference methods. These results demonstrate that coupling reinforcement learning with foundation models enables a paradigm shift from static embedding analysis to dynamic trajectory modeling, providing a powerful engine for discovering sequential perturbation strategies in regenerative medicine.

## Introduction

The advent of single-cell RNA sequencing (scRNA-seq) has revolutionized our understanding of cellular differentiation by revealing the heterogeneity and continuous nature of cell state transitions. ScRNA-seq captures intermediate differentiation states and enables the reconstruction of developmental trajectories at unprecedented resolution. This technology has revealed that differentiation is not a series of discrete steps but rather a continuous process through high-dimensional gene expression space^1–3^.

Recent advances in machine learning have led to the development of single-cell foundation models. Models such as scFoundation, Geneformer, scGPT, and scBERT learn rich representations of cellular states by encoding gene expression patterns into high-dimensional embeddings^4–7^. These models capture complex gene-gene relationships through attention mechanisms, enabling them to predict the effects of genetic perturbations without requiring prior knowledge of regulatory networks. However, current single-cell foundation models possess a critical limitation: they excel at predicting the effect of individual perturbations but lack the capacity for multi-step trajectory planning. Cellular differentiation is inherently a sequential process requiring coordinated regulation across multiple stages^8^. Identifying optimal perturbation sequences is essential for efficient protocol design, yet this problem is computationally intractable through exhaustive search given the combinatorial explosion of possible multi-step interventions.

Reinforcement learning (RL) has emerged as a powerful paradigm for solving sequential decision-making problems in complex environments. RL agents learn optimal strategies by interacting with environments, receiving rewards for beneficial actions, and iteratively updating their policies to maximize cumulative rewards. This approach has achieved remarkable success across diverse domains, from game playing and autonomous systems to robotic control ^9–12^. Recent applications have extended RL to molecular design and treatment strategy optimization^13,14^, demonstrating potential for biological problems.

In reinforcement learning, a world model is an internal representation of the environment that enables agents to predict future states and simulate potential action sequences before committing to real-world interventions. Single-cell foundation models function analogously as biological world models: trained on millions of cellular profiles, they compress complex signals into compact embeddings that capture cellular identity and differentiation potential. Through in silico perturbations, these models can “imagine” alternative cell states, predicting how genetic changes alter cellular phenotypes without experimental validation. However, current applications remain limited to single-step predictions, while cellular differentiation requires coordinated multi-step regulation across developmental stages. Identifying optimal perturbation sequences poses a combinatorial challenge intractable through exhaustive search. Integrating single-cell foundation models as differentiable simulators with reinforcement learning offers a natural solution: RL agents can learn to navigate complex differentiation landscapes through sequential decision-making, selecting actions that maximize long-term differentiation objectives. Induced pluripotent stem cell (iPSC) differentiation provides an ideal proof-of-concept for this integration.

Induced pluripotent stem cells (iPSCs) represent one of the most significant breakthroughs in regenerative medicine, offering unprecedented opportunities to model disease mechanisms, discover therapeutic compounds, and develop cell replacement therapies. Significant technical barriers limit the clinical application of iPSC technology. Current reprogramming and differentiation protocols are inefficient, time-consuming, and often yield heterogeneous cell populations with incomplete differentiation^15^. The presence of undifferentiated cells poses a critical safety risk, as these cells retain the capacity for unlimited self-renewal and can form teratomas upon transplantation^16^. Furthermore, existing protocols rely heavily on empirical optimization and prior biological knowledge, making them difficult to adapt across cell types, patient backgrounds, and culture conditions. Rational, data-driven approaches to protocol design could address these limitations.

The application of RL to cellular differentiation protocol design remains largely unexplored. Existing computational approaches to protocol optimization face several fundamental limitations: they either provide static predictions without sequential optimization capability, require explicit construction of regulatory networks, or operate on simplified representations that fail to capture cellular complexity^17–21^. No current approach systematically identifies optimal sequences of interventions needed to guide cells efficiently through complex differentiation trajectories while avoiding unwanted differentiation paths.

Early computational efforts to optimize protocols relied on bulk RNA sequencing and chromatin accessibility profiling to identify key regulatory factors. Methods such as CellNet and IRENE have been developed to reconstruct gene regulatory networks from bulk transcriptomic data, enabling researchers to predict the effects of transcription factor perturbations on cell identity^20,22^. However, bulk profiling approaches mask cellular heterogeneity and fail to capture the dynamic, intermediate states that characterize differentiation processes. Recent method like CellOracle has leveraged single-cell data to construct gene regulatory networks for in silico perturbation analysis^17^. While these approaches represent significant advances, they remain constrained to single-step predictions and require explicit construction of regulatory networks.

The application of RL to cellular differentiation protocol design remains largely unexplored, and no current approach systematically combines the rich cellular representations learned by foundation models with the sequential optimization capabilities of reinforcement learning. Bridging this gap could enable computational agents to discover optimal multi-step perturbation strategies that guide cells efficiently through complex differentiation trajectories.

Here, we introduce Differentiation with Reinforcement Learning (DiRL), a reinforcement learning framework that leverages single-cell foundation models as biological world models to identify optimal sequential genetic perturbations for cellular differentiation. We formulate differentiation protocol design as a goal-conditioned sequential decision-making problem: given a starting cell state and target cell type, an RL agent learns to select actions (genetic perturbations) that maximize differentiation efficiency. Foundation models serve as simulators that predict the cellular consequences of perturbations, enabling the RL agent to explore vast action spaces efficiently.

Our framework makes several key contributions. We develop a goal-conditioned actor-critic architecture that learns cell-type-specific differentiation policies, enabling a single model to guide cells toward diverse lineages. The framework is architecture-agnostic, demonstrating applicability across multiple foundation models and positioning it to leverage future advances in foundation model development. Importantly, learned policies recapitulate known differentiation mechanisms and critic value functions provide biologically meaningful orderings comparable to established pseudotime methods, validating that the RL framework captures genuine biological dynamics rather than exploiting model artifacts.

## Results

### Goal-Conditioned RL Agents Outperform Random Perturbation Baselines

We formulate differentiation as a sequential decision-making problem amenable to reinforcement learning and implement DiRL (**Figure 1**). To incentive RL for navigating cellular landscape toward target cell states, we derive rewards from cosine similarities of cell embeddings, where cell embeddings close to target cell states are assigned high rewards (**Methods**). Compared to using cosine similarities as rewards, our reward functions provide more robust and stable signals to guide RL agents for making optimal perturbation steps.

**Figure 1.**
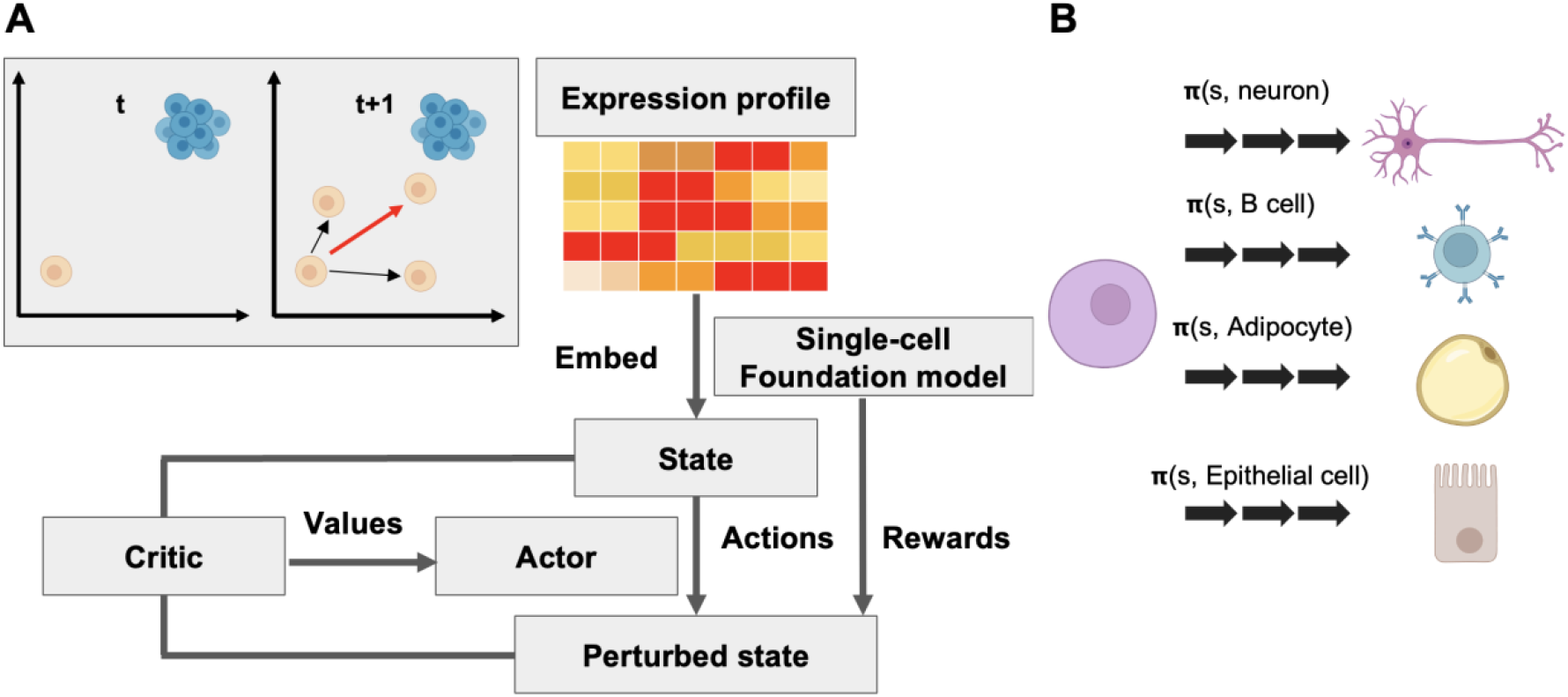
Overview of Differentiation with Reinforcement Learning (DiRL). (A) Framework schematic. DiRL navigates the cellular latent space encoded by single-cell foundation models to identify sequential genetic perturbations that drive transitions toward target cell states. The agent is trained to maximize a cosine-similarity-based reward. The Critic models the reward landscape using the underlying foundation model, while the Actor optimizes the perturbation policy based on feedback from the Critic. (B) Goal-conditioned learning. The DiRL agent learns a versatile policy capable of directing stem cells toward diverse terminal cell types.

Our approach employs a goal-conditioned actor-critic architecture. The actor network takes current cellular states and target cell types as input, outputting actions (genetic perturbations). The critic network evaluates action quality based on progress toward target cell states. This framework enables learning cell-type-specific policies that can navigate the complex landscape of cellular differentiation trajectories.

We implemented DiRL on three widely-used single-cell foundation models: scGPT, Geneformer, and STATE-embedding^6,7,23^, to assess generalizability across diverse architectures. These models employ distinct encoding strategies: Geneformer represents cells as ranked gene lists, while scGPT and STATE use binned expression values with transformer architectures.

To establish whether reinforcement learning can effectively learn trajectories leading to target cell states, we compared rewards obtained from DiRL to those obtained from random perturbation baseline. We evaluated DiRL on human intestinal organoids scRNA-seq dataset^24^ (n=64,603 cells spanning 19 cell types). DiRL was trained on the dataset and used for predicting trajectories toward various goals. To enable an unbiased benchmark, the same dataset was used as starting cell states and random trajectories of matching horizons were generated by uniformly sampling genes and expression changes at each decision step (**Methods**).

To compare DiRL-generated trajectories with random trajectories, we used win rate (proportion of trajectories with higher averaged rewards in model prediction compared to random baselines, see **Methods**). Across all three foundation model architectures, RL-trained policies based on scGPT and Geneformer significantly outperformed random perturbation baselines (**Figure 2A**). The scGPT-based agent achieved the highest performance, with a 69% win rate. Geneformer-based agents achieved a 54% win rate, demonstrating significant but modest improvements over random sampling. Notably, STATE-based agents achieved only 38% win, performing worse than the baseline.

**Figure 2.**
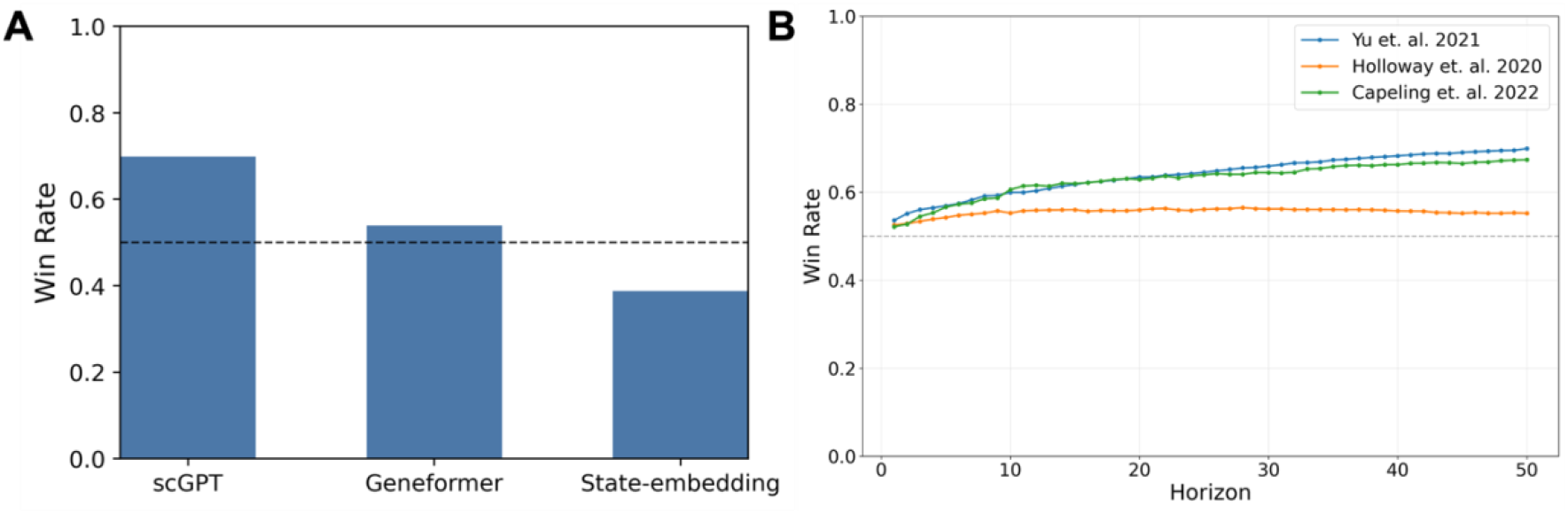
Benchmarking DiRL performance. (A) Foundation model comparison. scGPT, Geneformer, and STATE were evaluated as backbone for the DiRL framework. Win rates are defined as the percentage of trials where DiRL agents achieved higher final rewards than random perturbation baselines. (B) Cross-dataset generalization and scalability. scGPT-based models were benchmarked to assess robustness across varying trajectory lengths. Models were trained with a horizon of 10 and evaluated on a horizon of 50. Win rates increased with horizon, indicating that the learned policy effectively generalizes to longer differentiation trajectories.

### Performance Scales with Planning Horizon and Generalizes Across Datasets

To assess cross-dataset generalization, we applied the scGPT-based DiRL to three independent organoid datasets^24–26^. The datasets include organoids differentiated from both human embryonic stem cells and induced pluripotent stem cells using established directed differentiation protocols. Collectively, these datasets model early human intestinal development and contain diverse lineages. Despite differences in differentiation protocols and sources of human pluripotent stem cells, DiRL consistently outperformed random baselines across all datasets with win rates of 69.9%, 55.2%, and 67.4% respectively.

A key advantage of reinforcement learning over single-step prediction methods is the capacity for long-horizon planning. To evaluate whether DiRL genuinely perform multi-step reasoning rather than simply chaining independent one-step predictions, we systematically varied planning horizon length and measured performance scaling. Using the scGPT-based agent, we evaluated trajectories with horizons H steps from 1 to 50 across different datasets. Win rates over random baselines increased systematically with longer planning horizons (**Figure 2B**). This demonstrates that longer planning horizons enable more sophisticated navigation of the differentiation landscape, reaching higher-quality cell states through multi-step interventions. The consistent above-random performance across all contexts demonstrates the scalability of our framework to longer horizons.

### Learned Policies Prioritize Biologically Relevant Differentiation Regulators

While performance metrics establish that DiRL effectively guide differentiation trajectories, a critical question remains: do learned policies recapitulate known biological mechanisms, or do they exploit artifacts of foundation model predictions? To address this, we analyzed the genes selected by scGPT-based DiRL conditioned on diverse goal cell types and assessed their biological relevance to differentiation processes.

Actor-selected genes exhibited strong cell type specificity and were enriched for transcription factors and signaling proteins known to regulate developmental pathways (**Figure 3A**). The agent prioritized well-established master regulators of cell fate specification, including LHX1^27^ (critical for mesendoderm development), GLI1^28^ (key effector of Hedgehog signaling governing neural patterning), and NKX6-2^29^ (regulator of central nervous system development). Beyond transcription factors, the policy selected context-appropriate pathway modulators: Wnt signaling components (LEF1, LGR5, and the Wnt inhibitor FRZB), secreted growth factors (BMP2^30,31^ from the TGF-β family for dorsoventral patterning), and neuropeptides involved in neuronal maturation (VGF). The coherent selection of functionally related regulators demonstrates that DiRL learns biologically interpretable policies that recapitulate known differentiation mechanisms.

**Figure 3.**
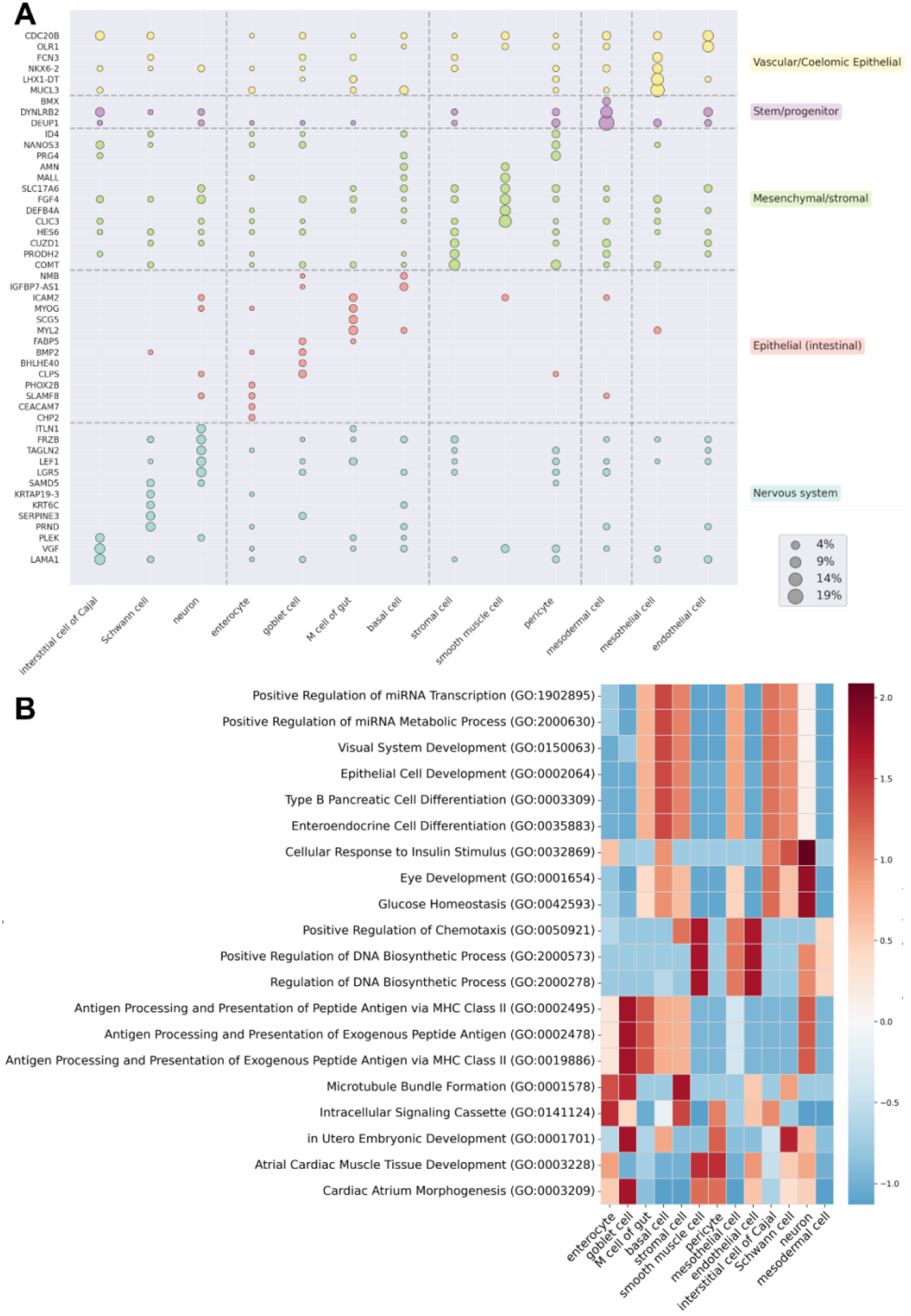
DiRL identifies cell fate regulators. (A) Goal-specific gene selection. Dot size represents the selection frequency of specific genes for each goal cell type. The distinct distribution of actions indicates that DiRL learns lineage-specific control policies. (B) Pathway-level regulatory scores. Summary scores represent the net regulatory activity, calculated by subtracting suppression frequency from activation frequency, normalized by pathway size and total action counts. Scores were z-score normalized across goal cell types; high z-scores indicate a high degree of specific activation for the target lineage.

### Critic Value Functions Provide Biologically Meaningful Pseudotime Orderings

The critic network estimates expected future rewards. Intuitively, critic values can serve as a proxy for differentiation progress. We first validated this by correlating critic values with experimental time points (**Figure 4A**). The learned critic values demonstrated strong temporal alignment, significantly outperforming random baselines and achieving correlation strength comparable to Palantir pseudotime. To further quantify biological fidelity, we assessed whether the critic captures the underlying gene programs driving differentiation. We identified genes correlated with critic values and calculated their overlap with ground-truth differentially expressed genes (DEGs) defined between stem and target cells. Critic-based orderings recovered DEGs at rates significantly exceeding random permutations and comparable to Palantir (**Figure 4B**). These quantitative metrics were reflected in lineage-consistent expression patterns (**Figure 4C**). High-value cell states were enriched for established cell type markers. According to PanglaoDB^32^, several top correlated genes are established cell type markers: CD24, LGALS4, MUC13, TSPAN1, and KRT18 for epithelial cells; ETS1 for endothelial cells; COL1A2 for mesenchymal cells; MDK for stromal cells; STMN1 for neural progenitors; and PFN1 for neurons.

**Figure 4.**
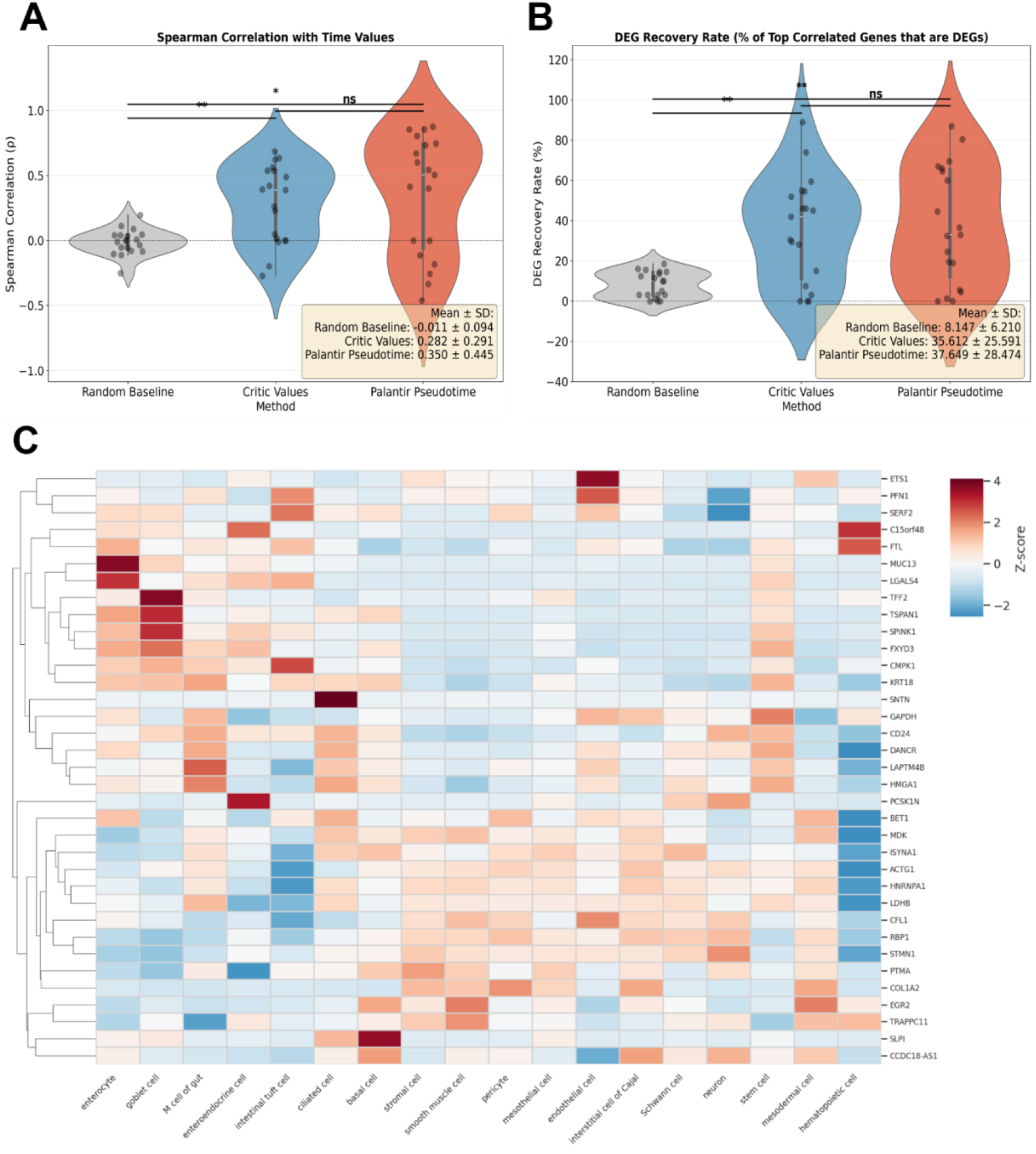
Critic values recapitulate gene expression dynamics during differentiation. (A) Temporal consistency. Learned critic values were compared against experimental time points (days in culture) from the intestinal organoid dataset to evaluate alignment with physical developmental time. (B) Biological validation of critic ordering. Genes correlated with either critic values or Palantir pseudotime were assessed for overlap with ground-truth differentially expressed genes (DEGs) for each goal cell type. Critic-based orderings recovered DEGs at rates comparable to Palantir and significantly superior to random baselines (generated by permuting cell order). (C) Top correlated genes exhibited lineage-consistent expression patterns (Two-tailed Mann–Whitney test; **: p < 0.01, *: p < 0.05).

Together, these results demonstrate that critic value functions encode biologically meaningful developmental information comparable to dedicated trajectory inference methods. This suggests that the RL framework captures genuine cell fate dynamics rather than spurious correlations, and that critic values may serve as interpretable metrics for assessing differentiation progression.

## Discussion

We developed DiRL, a reinforcement learning framework that leverages single-cell foundation models to identify optimal multi-step genetic perturbation sequences for cellular differentiation. Applied to organoid datasets, DiRL significantly outperformed random perturbation baselines and demonstrated scalability with planning horizon and biological interpretability through gene selection analysis and critic-based pseudotime ordering. These findings establish reinforcement learning as a viable paradigm for computationally guided differentiation.

DiRL uniquely combines several advantages: (1) multi-step trajectory optimization through reinforcement learning, (2) model training does not require explicit GRN construction, (3) goal-conditioned architecture enabling simultaneous learning across multiple target cell types, and (4) scalability to long planning horizons. The foundation model-based world model enables efficient exploration of vast perturbation spaces without exhaustive search.

Among the tested architectures, scGPT-based agents substantially outperformed Geneformer and STATE. scGPT likely outperforms competitors because its value-binned encoding retains absolute expression magnitudes, facilitating the precise state tracking necessary for sequential prediction. In contrast, Geneformer’s rank-based encoding improves robustness but discards abundance data, limiting the agent’s ability to leverage fine-grained perturbations. Finally, STATE’s reliance on population-level objectives appears to obscure the single-cell transition logic required for modeling perturbations. These findings demonstrate that reinforcement learning can effectively leverage foundation model predictions to guide differentiation trajectories, but performance depends on design of the underlying foundation model.

Several limitations warrant consideration. First, our framework operates on 2,000 highly variable genes selected for each dataset. While this captures major transcriptional variation and enables tractable action spaces, it may miss rare cell-type-specific regulators or context-dependent factors outside the selected gene set. Future work should explore adaptive gene selection strategies or hierarchical action spaces that first select regulatory modules, then specific genes. Second, we model differentiation as a Markov Decision Process (MDP), assuming future states depend only on current states and actions. However, cellular differentiation involves epigenetic memory, stochastic gene expression dynamics, and potentially non-Markovian dependencies where full trajectory history influences future states. Sequence-aware architectures such as transformer-based world models, may better capture trajectory-dependent dynamics and improve long-horizon planning^33,34^.

Efficient and reliable differentiation protocols are foundational for regenerative medicine. Current protocol development requires empirical optimization, limiting the pace of clinical translation. DiRL offers a complementary computational approach that could accelerate this process by systematically exploring vast perturbation spaces, identifying promising candidates for experimental validation, and revealing unexpected regulatory strategies.

From a basic science perspective, analyzing learned policies provides insights into cell fate control. Genes consistently selected across diverse goals may represent universal differentiation regulators, while goal-specific genes reveal lineage-determining factors. Furthermore, the critic value functions offer an interpretable metric for quantifying differentiation progression. These value estimates provide a comparable, dynamics-based alternative for ordering cellular states. Ultimately, this highlights the broader potential of single-cell foundation models to move beyond static representation learning, acting instead as predictive engines for capturing and directing cell state transitions.

## Methods

### Goal-Conditioned Reinforcement Learning Formulation

We model the cellular differentiation process as a Markov Decision Process (MDP) with trajectories 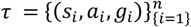, where *s*_*i*_ represents cell states (foundation model embeddings), *a*_*i*_ denotes actions (gene perturbations), and *g*_*i*_ indicates target cell types (goals). Actions are factorized as *a*_*t*_ = (*a*_*gene*_, *a*_*expr*_), where *a*_*gene*_ selects the gene to perturb and *a*_*expr*_ determines the expression change magnitude.

We adopted actor-critic architecture. The actor learns a policy *π*_*θ*_(*a*|*s, g*) that maximizes expected cumulative rewards, while the critic estimates the expected future rewards *V*_*φ*_(*s,g*) given the current state and goal. To ensure efficient learning, we restrict goal sampling within biologically relevant clusters identified using Leiden clustering (resolution = 1.0, neighbors = 10).

### Reward Function Design

Given a current cell state *s*, target cell type 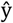, and perturbed state 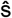(the state after applying action *a*_*t*_), the total reward is: *r*_*t*_ = *R*_*goal*_ + *R*_*potential*_ + *R*_*prob*_

The goal achievement reward provides a binary completion signal: 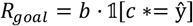 where *b* is the reward magnitude, *c*∗ is the best matching cell type (defined below), and 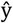 is the target cell type.

### Potential-Based Reward Shaping

To provide intermediate guidance, we implement potential-based reward shaping. The potential function *Φ*(*s*) measures the average similarity to all cells of the target type within the goal cluster:

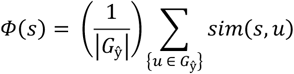

where *G*_ŷ_ represents all cells of target type ŷ in the cluster, and *sim*(·,·) is the cosine similarity function. The potential reward encourages progress toward the goal: *Rpotential* = *λ*(*γ* · *Φ*(ŝ) − *Φ*(*s*)) where *λ* is the potential reward coefficient and *γ* is the discount factor.

### Probability-Based Reward

We implement a cluster-based soft matching mechanism using rank-decayed weighting. For cluster embeddings {*e*_*j*_} with cell type labels {*y*_*j*_}, we compute distances *d*_*j*_ between the perturbed state ŝ and each cluster embedding, then derive similarities and ranks. The rank-decayed weights are: *w*_*j*_ = exp(−*α* · *rank*_*j*_), where *α* = 0.01. The probability of belonging to cell type *c* given state ŝ is:

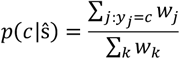

The best matching cluster is determined as: *c∗* = *argmaxcp*(*c*|ŝ). The probability reward directly measures the likelihood of reaching the target cell type ŷ: *Rprob* = *p*(ŷ|ŝ). This unified framework ensures that goal achievement is triggered when the most likely cell type *c∗* matches the target ŷ, while the *Rprob* provides continuous feedback about progress toward the target.

### Policy Optimization with Proximal Policy Optimization (PPO)

We employ PPO to ensure stable policy updates by constraining the policy change magnitude. PPO maintains a reference policy *π*_*old*_ (the policy from the previous training iteration) and computes the probability ratio:

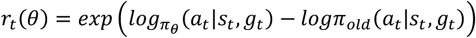

where *θ* represents the policy parameters, *s*_*t*_, *a*_*t*_, and *g*_*t*_ are the state, action, and goal at time *t* respectively.

The clipped policy loss prevents destructive policy updates:

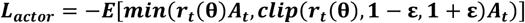

where *A*_*t*_ represents the advantage estimate, *ε* is the clipping parameter (default to 0.2), and the expectation is taken over the batch of experiences.

### Value Function Learning and Entropy Regularization

The critic network learns to estimate the expected cumulative reward using mean squared error loss:

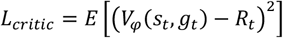

where *φ* represents the value function parameters and *R*_*t* is the empirical return.

To maintain exploration, we add entropy regularization, *H* = *E*[*H*(*π*_*θ*_(· |*s*_*t*_, *g*_*t*_))]. The combined objective function is:

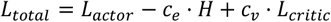

where *c*_*e*_ = 0.001 and *c*_*v*_ = 0.5.

### Advantage Estimation and Return Computation

We employ Generalized Advantage Estimation (GAE) to compute low-variance advantage estimates that balance bias and variance in policy gradient estimation. The temporal difference (TD) error is: *δ*_*t*_ = *r*_*t*_ + *γ*(1 − *d*_*t*_)*V*(*s*_*t*_+1, *g*_*t*_+1) − *V*(*s*_*t*_, *g*_*t*_) where γ is the discount factor and *d*_*t*_ indicates episode termination.

GAE computes advantages recursively:

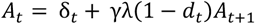

Returns combine advantages with value estimates:

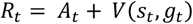

### Action Space Factorization

Actions are factorized into gene selection and expression modification:

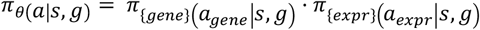

Both components use categorical distributions with log probabilities:

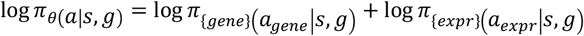

We adopt Geneformer, scGPT, and STATE-embedding as simulator. For Geneformer, gene expression matrices are represented as ranked gene lists. Expression changes *a*_*expr*_ represent relative position displacements (range: -10 to +10, scaled by 100). Absent genes are inserted at the center position before displacement. For scGPT and STATE-embedding, perturbations modify expression bins of selected genes. If the selected gene is absent, a randomly chosen gene is replaced.

### Benchmark

To benchmark model performance, we generated random baselines using uniform sampling from available genes and expression changes. Trajectories were compared with win rate based on mean probability rewards, providing a robust evaluation metric for differentiation protocol effectiveness. Win-rate: 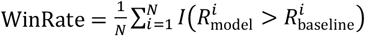 where *N* is the total number of episodes, *I* is the indicator function, 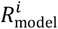 is the average probability rewards obtained from trajectory *i* generated by trained models, and 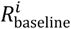 is the average rewards obtained from baseline at the same

### Critic-value analysis

We validated the biological relevance of the learned critic function using two metrics. First, we calculated the correlation between single-cell critic values and experimental time points to assess temporal consistency. Second, we evaluated the recovery of gene expression dynamics. We ordered cells by their critic values and identified genes differentially expressed between low-critic-value (early-stage) and high-critic-value (late-stage) cells. We then calculated the overlap between these critic-derived genes and a ground-truth set of DEGs computed between the starting stem cells and terminally differentiated target cells. Performance was benchmarked against Palantir pseudotime and randomized cell orderings.

## Code Availability

All code necessary to replicate DiRL, including the implementation of the Actor-Critic networks, training protocols, and the scripts used for benchmarking scGPT, Geneformer, and STATE foundation models are available at https://github.com/KaiLiChang/DiRL.

## Acknowledgments

This study was supported by the Cancer Prevention and Research Institute of Texas (CPRIT) award RP240131.

## References

1. Joost, S., Zeisel, A., Jacob, T., Sun, X., La Manno, G., Lönnerberg, P., Linnarsson, S., and Kasper, M. (2016). Single-Cell Transcriptomics Reveals that Differentiation and Spatial Signatures Shape Epidermal and Hair Follicle Heterogeneity. cels 3, 221-237.e9. 10.1016/j.cels.2016.08.010.

2. Dulken, B.W., Leeman, D.S., Boutet, S.C., Hebestreit, K., and Brunet, A. (2017). Single-Cell Transcriptomic Analysis Defines Heterogeneity and Transcriptional Dynamics in the Adult Neural Stem Cell Lineage. Cell Reports 18, 777–790. 10.1016/j.celrep.2016.12.060.

3. Pellin, D., Loperfido, M., Baricordi, C., Wolock, S.L., Montepeloso, A., Weinberg, O.K., Biffi, A., Klein, A.M., and Biasco, L. (2019). A comprehensive single cell transcriptional landscape of human hematopoietic progenitors. Nat Commun 10, 2395. 10.1038/s41467-019-10291-0.

4. Hao, M., Gong, J., Zeng, X., Liu, C., Guo, Y., Cheng, X., Wang, T., Ma, J., Zhang, X., and Song, L. (2024). Large-scale foundation model on single-cell transcriptomics. Nat Methods, 1–11. 10.1038/s41592-024-02305-7.

5. Yang, F., Wang, W., Wang, F., Fang, Y., Tang, D., Huang, J., Lu, H., and Yao, J. (2022). scBERT as a large-scale pretrained deep language model for cell type annotation of single-cell RNA-seq data. Nat Mach Intell 4, 852–866. 10.1038/s42256-022-00534-z.

6. Cui, H., Wang, C., Maan, H., Pang, K., Luo, F., and Wang, B. (2023). scGPT: Towards Building a Foundation Model for Single-Cell Multi-omics Using Generative AI. Preprint, 10.1101/2023.04.30.538439.

7. Theodoris, C.V., Xiao, L., Chopra, A., Chaffin, M.D., Al Sayed, Z.R., Hill, M.C., Mantineo, H., Brydon, E.M., Zeng, Z., Liu, X.S., et al. (2023). Transfer learning enables predictions in network biology. Nature 618, 616–624. 10.1038/s41586-023-06139-9.

8. Cerneckis, J., Cai, H., and Shi, Y. (2024). Induced pluripotent stem cells (iPSCs): molecular mechanisms of induction and applications. Sig Transduct Target Ther 9, 112. 10.1038/s41392-024-01809-0.

9. Silver, D., Huang, A., Maddison, C.J., Guez, A., Sifre, L., van den Driessche, G., Schrittwieser, J., Antonoglou, I., Panneershelvam, V., Lanctot, M., et al. (2016). Mastering the game of Go with deep neural networks and tree search. Nature 529, 484–489. 10.1038/nature16961.

10. OpenAI, Andrychowicz M., Baker, B., Chociej, M., Jozefowicz, R., McGrew, B., Pachocki, J., Petron, A., Plappert, M., Powell, G., et al. (2019). Learning Dexterous In-Hand Manipulation. Preprint at arXiv, 10.48550/arXiv.1808.00177.

11. Shao, K., Tang, Z., Zhu, Y., Li, N., and Zhao, D. (2019). A Survey of Deep Reinforcement Learning in Video Games. Preprint at arXiv, 10.48550/arXiv.1912.10944.

12. Sallab, A.E., Abdou, M., Perot, E., and Yogamani, S. (2017). Deep Reinforcement Learning framework for Autonomous Driving. ei 29, 70–76. 10.2352/ISSN.2470-1173.2017.19.AVM-023.

13. Popova, M., Isayev, O., and Tropsha, A. (2018). Deep reinforcement learning for de novo drug design. Science Advances 4, eaap7885. 10.1126/sciadv.aap7885.

14. Komorowski, M., Celi, L.A., Badawi, O., Gordon, A.C., and Faisal, A.A. (2018). The Artificial Intelligence Clinician learns optimal treatment strategies for sepsis in intensive care. Nat Med 24, 1716–1720. 10.1038/s41591-018-0213-5.

15. Cerneckis, J., Cai, H., and Shi, Y. (2024). Induced pluripotent stem cells (iPSCs): molecular mechanisms of induction and applications. Sig Transduct Target Ther 9, 112. 10.1038/s41392-024-01809-0.

16. Yamanaka, S. (2020). Pluripotent Stem Cell-Based Cell Therapy—Promise and Challenges. Cell Stem Cell 27, 523–531. 10.1016/j.stem.2020.09.014.

17. Kamimoto, K., Stringa, B., Hoffmann, C.M., Jindal, K., Solnica-Krezel, L., and Morris, S.A. (2023). Dissecting cell identity via network inference and in silico gene perturbation. Nature 614, 742–751. 10.1038/s41586-022-05688-9.

18. Li, C., Chen, S., Chen, Y., Bian, H., Hao, M., Wei, L., and Zhang, X. (2024). scDirect: key transcription factor identification for directing cell state transitions based on single-cell multi-omics data. Preprint at bioRxiv, 10.1101/2024.01.08.574757.

19. Hammelman, J., Patel, T., Closser, M., Wichterle, H., and Gifford, D. (2022). Ranking reprogramming factors for cell differentiation. Nat Methods 19, 812–822. 10.1038/s41592-022-01522-2.

20. Jung, S., Appleton, E., Ali, M., Church, G.M., and del Sol, A. (2021). A computer-guided design tool to increase the efficiency of cellular conversions. Nat Commun 12, 1659. 10.1038/s41467-021-21801-4.

21. Cahan, P., Li, H., Morris, S.A., Lummertz da Rocha, E., Daley, G.Q., and Collins, J.J. (2014). CellNet: Network Biology Applied to Stem Cell Engineering. Cell 158, 903–915. 10.1016/j.cell.2014.07.020.

22. Cahan, P., Li, H., Morris, S.A., da Rocha, E.L., Daley, G.Q., and Collins, J.J. (2014). CellNet: Network Biology Applied to Stem Cell Engineering. Cell 158, 903–915. 10.1016/j.cell.2014.07.020.

23. Adduri, A.K., Gautam, D., Bevilacqua, B., Imran, A., Shah, R., Naghipourfar, M., Teyssier, N., Ilango, R., Nagaraj, S., Dong, M., et al. (2025). Predicting cellular responses to perturbation across diverse contexts with State. Preprint at bioRxiv, 10.1101/2025.06.26.661135.

24. Xu, Q., Halle, L., Hediyeh-zadeh, S., Kuijs, M., Riedweg, R., Kilik, U., Recaldin, T., Yu, Q., Rall, I., Frum, T., et al. (2025). An integrated transcriptomic cell atlas of human endoderm-derived organoids. Nat Genet 57, 1201–1212. 10.1038/s41588-025-02182-6.

25. Yu, Q., Kilik, U., Holloway, E.M., Tsai, Y.-H., Harmel, C., Wu, A., Wu, J.H., Czerwinski, M., Childs, C.J., He, Z., et al. (2021). Charting human development using a multi-endodermal organ atlas and organoid models. Cell 184, 3281-3298.e22. 10.1016/j.cell.2021.04.028.

26. Holloway, E.M., Wu, J.H., Czerwinski, M., Sweet, C.W., Wu, A., Tsai, Y.-H., Huang, S., Stoddard, A.E., Capeling, M.M., Glass, I., et al. (2020). Differentiation of Human Intestinal Organoids with Endogenous Vascular Endothelial Cells. Developmental Cell 54, 516-528.e7. 10.1016/j.devcel.2020.07.023.

27. Zhao, Y., Kwan, K.-M., Mailloux, C.M., Lee, W.-K., Grinberg, A., Wurst, W., Behringer, R.R., and Westphal, H. (2007). LIM-homeodomain proteins Lhx1 and Lhx5, and their cofactor Ldb1, control Purkinje cell differentiation in the developing cerebellum. Proceedings of the National Academy of Sciences 104, 13182–13186. 10.1073/pnas.0705464104.

28. Rowton, M., Perez-Cervantes, C., Hur, S., Jacobs-Li, J., Lu, E., Deng, N., Guzzetta, A., Hoffmann, A.D., Stocker, M., Steimle, J.D., et al. (2022). Hedgehog signaling activates a mammalian heterochronic gene regulatory network controlling differentiation timing across lineages. Dev Cell 57, 2181-2203.e9. 10.1016/j.devcel.2022.08.009.

29. Dorboz, I., Aiello, C., Simons, C., Stone, R.T., Niceta, M., Elmaleh, M., Abuawad, M., Doummar, D., Bruselles, A., Wolf, N.I., et al. (2017). Biallelic mutations in the homeodomain of NKX6-2 underlie a severe hypomyelinating leukodystrophy. Brain 140, 2550–2556. 10.1093/brain/awx207.

30. Lillien, L., and Raphael, H. (2000). BMP and FGF regulate the development of EGF-responsive neural progenitor cells. Development 127, 4993–5005. 10.1242/dev.127.22.4993.

31. Meyers, E.A., and Kessler, J.A. (2017). TGF-β Family Signaling in Neural and Neuronal Differentiation, Development, and Function. Cold Spring Harb Perspect Biol 9, a022244. 10.1101/cshperspect.a022244.

32. Setty, M., Kiseliovas, V., Levine, J., Gayoso, A., Mazutis, L., and Pe’er, D. (2019). Characterization of cell fate probabilities in single-cell data with Palantir. Nat Biotechnol 37, 451–460. 10.1038/s41587-019-0068-4.

33. Robine, J., Höftmann, M., Uelwer, T., and Harmeling, S. (2023). Transformer-based World Models Are Happy With 100k Interactions. Preprint at arXiv, 10.48550/arXiv.2303.07109.

34. Chen, L., Lu, K., Rajeswaran, A., Lee, K., Grover, A., Laskin, M., Abbeel, P., Srinivas, A., and Mordatch, I. (2021). Decision Transformer: Reinforcement Learning via Sequence Modeling. Preprint at arXiv, 10.48550/arXiv.2106.01345.

